# How to make methodological decisions when inferring social networks

**DOI:** 10.1101/739789

**Authors:** André C. Ferreira, Rita Covas, Liliana R. Silva, Sandra C. Esteves, Inês F. Duarte, Rita Fortuna, Franck Theron, Claire Doutrelant, Damien R. Farine

## Abstract

Constructing and analysing social networks data can be challenging. When designing new studies, researchers are confronted with having to make decisions about how data are collected and networks are constructed, and the answers are not always straightforward. The current lack of guidance on building a social network for a new study system might lead researchers to try several different methods, and risk generating false results arising from multiple hypotheses testing. We suggest an approach for making decisions when developing a network without jeopardising the validity of future hypothesis tests. We argue that choosing the best edge definition for a network can be made using *a priori* knowledge of the species, and testing hypotheses that are known and independent from those that the network will ultimately be used to evaluate. We illustrate this approach by conducting a pilot study with the aim of identifying how to construct a social network for colonies of cooperatively breeding sociable weavers. We first identified two ways of collecting data using different numbers of feeders and three ways to define associations among birds. We then identified which combination of data collection and association definition maximised (i) the assortment of individuals into ‘breeding groups’ (birds that contribute towards the same nest and maintain cohesion when foraging), and (ii) socially differentiated relationships (more strong and weak relationships than expected by chance). Our approach highlights how existing knowledge about a system can be used to help navigate the myriad of methodological decisions about data collection and network inference.

**SIGNIFICANCE STATEMENT:** General guidance on how to analyse social networks has been provided in recent papers. However less attention has been given to system-specific methodological decisions when designing new studies, specifically on how data are collected, and how edge weights are defined from the collected data. This lack of guidance can lead researchers into being less critical about their study design and making arbitrary decisions or trying several different methods driven by a given preferred hypothesis of interest without realising the consequences of such approaches. Here we show that pilot studies combined with *a priori* knowledge of the study species’ social behaviour can greatly facilitate making methodological decisions. Furthermore, we empirically show that different decisions, even if data are collected under the same context (e.g. foraging), can affect the quality of a network.

## INTRODUCTION

Social network analysis (SNA) has gained popularity in behaviour ecology as a tool to study the processes underlying the associations between individuals and the consequences of those associations (Cantor et al. 2019). It allows biologists to characterize not only the social environment experienced by a single individual in the population, but also the broader social characteristics of a population (Newman 2010). However, while the methods involved in analysing a network are reasonably well-explained (e.g. Whitehead 2008), there are many decisions involved with the design of data collection and creating the network itself (Farine and Whitehead 2015). Decisions about the design of a study can have consequences on the inferred network structure. How can we know that our design decisions produce a meaningful network for the species and the type of hypotheses we are studying? There is generally little discussion of the considerations made when designing a network-based study, with most published papers presenting their design as a “fait accompli”.

When analysing a social network, the key decision that needs to be made is how to define the relationships (edges) connecting the individuals (nodes). This definition can include two main components. The first set of considerations relates to how data are collected, and the second relates to how observations are turned into edge weights. In most systems, the scope of decisions about data collection appears constrained by methodological limitations, but often there are choices that reflect some trade-offs. For example, is it better to collect fewer data across more individuals at once or to collect more detailed data on fewer individuals? Davis et al. (2018) provide useful discussion on the impact of these trade-offs. However, there is no general guidance on how to quantify the relative value of different approaches when faced with real data. Once data are collected, the second set of considerations that arise reflect decisions about how to calculate the strength of the relationships among individuals. While one aspect determining the accuracy of a network is collecting sufficient data (see Farine and Strandburg-Peshkin 2015), how data are used to generate quantitative measures of connection strength (edge weights) can also have a large impact on the resulting network. For example, different association indices (Cairns and Schwager 1987; Hoppitt and Farine 2018) or different resolutions of data (e.g. the number of grooming bouts vs. the amount of time spent grooming) can be used to estimate the strength of a given relationship.

The lack of guidance on how to evaluate a given data collection and network inference approach might lead researchers to try several different methods and to select the one that finds best correlates with the outcome they are studying (e.g. survival). Such a correlation could give a false impression that the method produces a network that is successfully capturing the species’ or population social structure. At worse, it could constitute a multiple hypotheses testing scenario, elevating rates of type I errors, especially when combined with opportunities to calculate multiple metrics (e.g. degree, betweenness, etc.) based on the network. For example, a researcher might be interested in understanding whether specific individual attributes, such as personality, correlates with one or multiple network centrality metrics (e.g. Aplin et al. 2013; Boogert et al. 2014a; Chock et al. 2017; Johnson et al. 2017; Moyers et al. 2018). In the absence of significant results, it could be tempting to change *a posteriori* the methods by which the network *is produced* using the same data, such as changing the time window or the proximity criterion used to consider that two individuals are associated. Hence, an important challenge arises when it is unknown whether failing to reject a hypothesis is a consequence of the expected pattern not being present or because the network was not correctly constructed. We therefore need an approach that avoids creating circularity, i.e. using the same data tested in different ways to corroborate a given hypotheses as well as using the significant result to corroborate the quality of the information contained in the network. This is particularly problematic because most published studies do not inform the readers about how decisions were made, such as whether they were made arbitrarily (or based on a published study), if they were based on pilot studies, or if they were explored in the way described above (but see: Castles et al. 2014; Boogert et al., 2014a; Mourier et al. 2017, for some exceptions).

Two approaches can help with making decisions about the design of a network study. The first is to collect pilot data, which is rarely feasible. The second is to run exploratory *a priori* analyses aimed at comparing different competing networks resulting from different data collection setups or network generation methods. Such a process can include testing and interpreting simple hypotheses that we generally consider a network from that species should support, *before* testing the hypothesis of interest. For example, in a species where mother and offspring create strong social bonds we expect that the implemented method would result in a network that would be able to capture these preferred associations (i.e. estimate the edge weight within a family as being significantly greater than those between other sets of individuals, see Boogert et al. 2014a). Such an analysis would then provide information about whether a network is capturing one or more important aspects of the biology of the system.

In this paper, we provide an empirical example of how to make decisions about the design of a network study using exploratory *a priori* test. We start by formulating simple hypotheses tests to help guide the design of data collection and network inference from a population of colonial and cooperatively breeding sociable weavers (*Philetairus socius*). In this population, individuals are individually marked with PIT-tags allowing automatic data collection at feeders containing supplemental food. Our work forms part of a broader study into the species’ social behaviour where we seek to understand whether specific individual attributes influences social relationships among the individuals within a colony. We decided to collect associations in a feeding context not only because this has been shown to be important and meaningful in other bird studies (e.g. Aplin et al. 2015a), but also as a result of the general insights on the social foraging behaviour of this species that have been reported in previous studies on this population (Rat et al. 2015; Lloyd et al. 2017; Silva et al., 2018). Therefore it seems reasonable to assume that information about social relationships within a colony could be obtained from foraging associations (see Farine 2015), if the study is well designed.

We evaluate the performance of different design decisions at extracting two fundamental structural aspect of the social system in our study species. The first metric is social differentiation, which we calculate using the coefficient of variation. Because sociable weavers’ colonies are large, we do not expect birds to have the same relationship strength with all colony members. Thus, an informative network should be one that features large differences in the connection strengths that individuals have in their social network (i.e. having many small and large values, rather than many intermediate values). However solely relying on social differentiation can be misleading as high values can be obtained as a result of non-social factors (e.g. low sampling or spatial distribution), nor does maximising social differentiation necessarily result in the most biologically accurate network. Thus, our second metric for testing if the edges in the foraging network reflect social bonds that transcend simple foraging synchrony is to calculate the assortment by breeding group. Sociable weaver colonies contain several breeding groups composed of breeders with their helpers (Covas et al. 2006). Assortment by breeding group captures the tendency of individuals from the same breeding group to be more strongly connected to one-another in the network. We expect this because while aggression between individuals at food patches is common (sociable weavers typically forage in large groups containing many colony members), aggression between members of the same breeding groups is rare (suggesting higher tolerance for other breeding group members Rat 2015). Thus, we expect members of the same breeding group to be disproportionately detected together, resulting in social networks that are assorted by breeding group membership.

First, we focus on the effects of different design decisions for data collection on the resulting values of social differentiation and assortment by breeding group. Specifically, we test the effects of allowing different numbers of individuals to feed simultaneously. This is an important decision for any researcher starting a new study as it can impact the robustness of the networks (Davis, et al. 2018). It is not clear whether sociable weavers with stronger social relationships feed more synchronously across repeated foraging visits than birds with weaker relationships, or whether the differences in behaviour are better defined as the patterns of foraging within a foraging visit (i.e. with who within the flock the individuals prefer to associate in close proximity). The former requires more widespread effort, while the latter requires more refined data to be collected within foraging flocks. We therefore compare different setups for collecting associations that differ in the number of birds that can be detected in an automated RFID system at the same time.

Second, we focus on how to define associations from within a given dataset. Specifically, we compare three different criteria to generate quantitative measures of edge weights in the network. Two criteria are based on number of co-occurrences in ‘gathering events’. These are akin to using the ‘gambit-of-the-group’ approach, where all birds that are detected (i.e. observed) in a flock together are considered to be associated. However, this approach discards more detailed data that could be available about within-flock structure, and instead assumes that birds with strong relationships will be co-observed in the same flock more often. The third approach is a more direct measure of the proportion of time that two individuals spend in close proximity within the flocks. That is, because we collected data at multiple readers in close proximity, we could estimate how much time two individuals spent on neighbouring feeders.

Our aim is to provide an example of the step-by-step procedure that can help guide researchers on what should be the right decisions for their network study. In doing so, our study also highlights how simple approaches, using short periods of pilot data collection and evaluations based on known factors about a study species, can facilitate making methodological decisions that could have long-term impact on the success of a study. Such an approach goes beyond studies using social network analysis.

## METHODS

### Study scope and model species

We studied a population of sociable weavers at Benfontein Nature Reserve, situated ca. 6 km south-east of Kimberley, in the Northern Cape Province, South Africa. The sociable weaver is endemic to the semi-arid savannahs of southern Africa (Maclean, 1973a) and feeds mainly on insects and seeds (Maclean, 1973c). Sociable weavers build large nests, usually on *Acacia* (*Vachellia*) trees, with several independent chambers where the birds roost throughout the year and where breeding takes place (Maclean, 1973b). This species exhibits three noticeable cooperative behaviours: building the communal nest, feeding nestlings of others, and communal nest defence from predators such as snakes (e.g. Boomslang, *Dyspholidus typus* and Cape cobra, *Naja nivea*). The size of a colony can range from less than ten to several hundred individuals. The breeding pairs can either breed with or without helpers (30-80% of breeding attempts have helpers; Covas et al. 2008).

This study is part of a long-term research programme which involves the annual capture of 14 colonies to maintain an individually marked population (all individuals are marked with a unique metal ring and colour combination: Covas et al. 2008; Paquet et al. 2015). At five colonies, all birds are also marked with a passive integrated transponder (PIT-tag, enclosed in a plastic leg ring). These ranged in size from 44 to 82 individuals (colony size estimated from the annual captures in September 2017). Additionally, blood samples are collected from all individuals for molecular determination of sex, parentage and relatedness among individuals (e.g. Covas et al. 2006; van Dijk et al. 2014).

### Breeding groups’ identification

Breeding groups were determined using video recordings of the chambers during the reproductive season of October 2017 to January 2018. We routinely inspected all colonies every 3 days to identify initiation of new clutches. We visited chambers in the days around the expected hatching date to determine the age of the nestlings and then recorded each breeding group for at least two hours when the chicks were between 8 - 20 days old. We considered an individual as part of the group if it was seen feeding the chicks at least 3 times, as occasionally some individuals try to feed but are expelled by the other members of the group.

### SNA data collection

During December 2017 and April 2018 we collected two rounds of association data in a feeding context using artificial feeders at the 5 PIT-tagged colonies. Data from a colony were collected at 2 feeding boxes (high density setup), each with 4 perches and 4 small standard plastic bird feeders. Each small feeder allowed for only one bird to feed at a time and was fitted with a RIFD antenna (Priority1rfid, Melbourne, Australia) connected to a data logger (Fig. 1a).

**Fig. 1.**
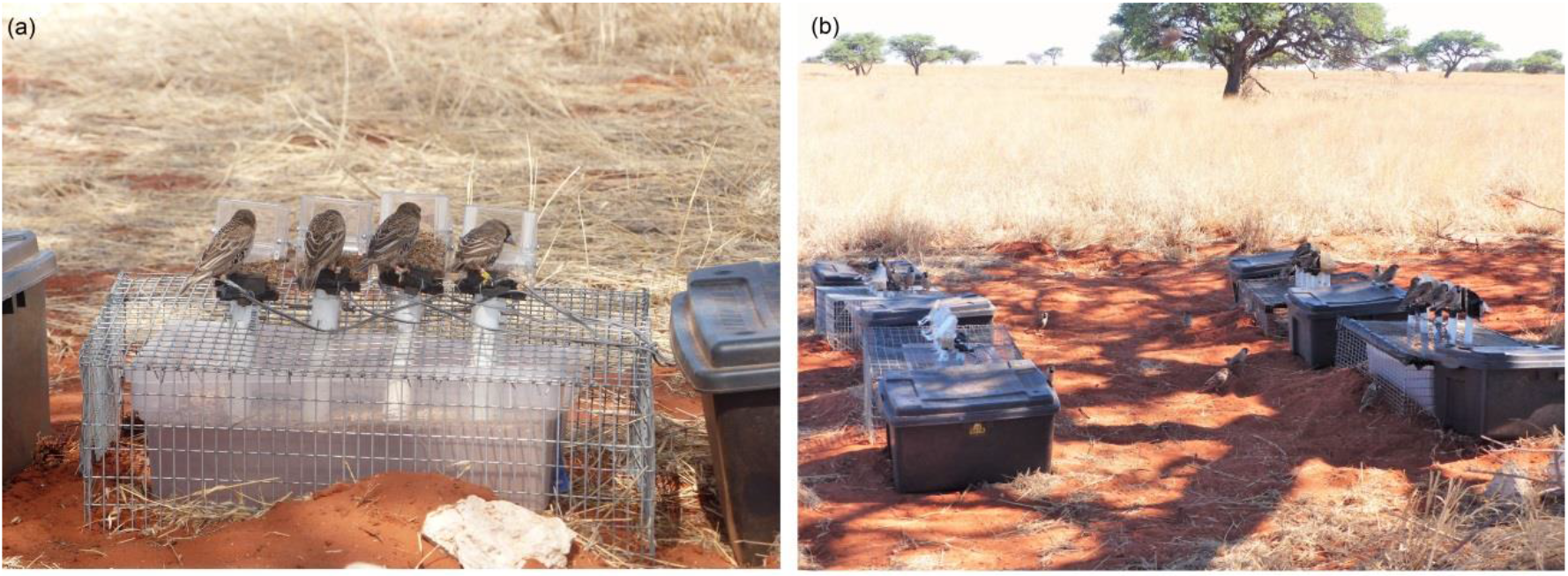
Setup for collecting associations (a) A feeding box with birds feeding at the four plastic feeders and the RFID antennas (b) the low density setup with four feeding boxes. Photographs by Cecile Vansteenberghe.

For two of the five colonies, feeding associations were also collected using a different setup that differed in the number of birds feeding at the same time. This setup featured 4 feeding boxes instead of 2 (low density setup; Fig. 1b), allowing birds to spread out more when visiting the feeding station (i.e. lower competition) and for us to collect more direct observations of co-feeding (more birds feeding simultaneously). Data for each setup were collected within the same time period, alternating each day for each setup (high or low density), in order to allow comparison of the two setups without a cofounding factor of time period in which the data were collected. We collected 10 days of data for each setup.

Data were collected for 14 days (sampled continuously) for each of the 3 colonies with one setup, and for 20 days in total for the other 2 colonies. For all the 5 colonies, the feeding location was 80-205 meters away from the colony.

Data from the 5 colonies allows for comparison of the different methods for edge weight calculations, while data from the 2 colonies collected with different number of feeding boxes also allow us to compare the networks from different data collection methods.

### Edge weight calculations

We calculated associations from our observation data in two different ways:

1. Co-occurrence method. We used the gambit of the group (see Whitehead and Dufault 1999; Franks et al. 2010), where all individuals that are observed together are considered to be equally connected to each other (i.e. a flock). There are several ways a flock can be defined (see Farine and Whitehead 2015). Here, we used an established method of inferring flocks based on the time differences between two detections. The start and end times of a ‘wave’ of individuals considered to be forming a flock are determined by a Gaussian mixture model (GMM; following Psorakis et al. 2015), which is an automated clustering algorithm designed to detect peaks in the temporal profile of activities at the artificial feeders. This approach uses data from the feeding behaviour of the entire set of individuals as part of determining the associations between any two individuals.
2. Time overlap method. We estimated association strengths using the total time that two individuals overlapped feeding at the same feeding box.

These two methods are described in more detail below. For the co-occurrence method two variants are used (see Fig. 2): one focused on the association at the broad flock level (single GMM) and the other based on association within the flock (double GMM). Therefore 3 different network types were compared for each combination of colony (see Fig. 3).

**Fig. 2.**
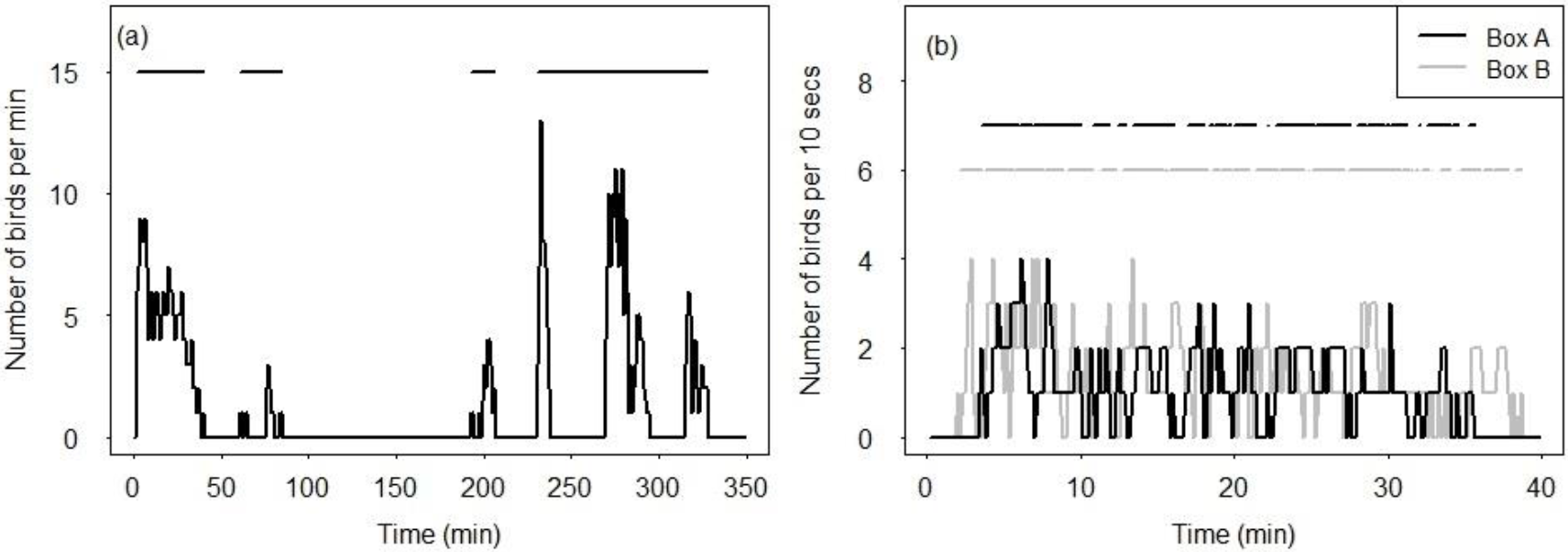
Example of applying the GMM algorithm method. (a) Sociable weaver visits to a feeding location during one morning. The top straight lines represent the gathering events resulting from the first GMM. (b) The gathering events resulting from the second GMM, discriminating between the two feeding boxes and using only visits from the first event determined by the first GMM (corresponding to the first horizontal line segment on Fig 2a).

**Fig. 3.**
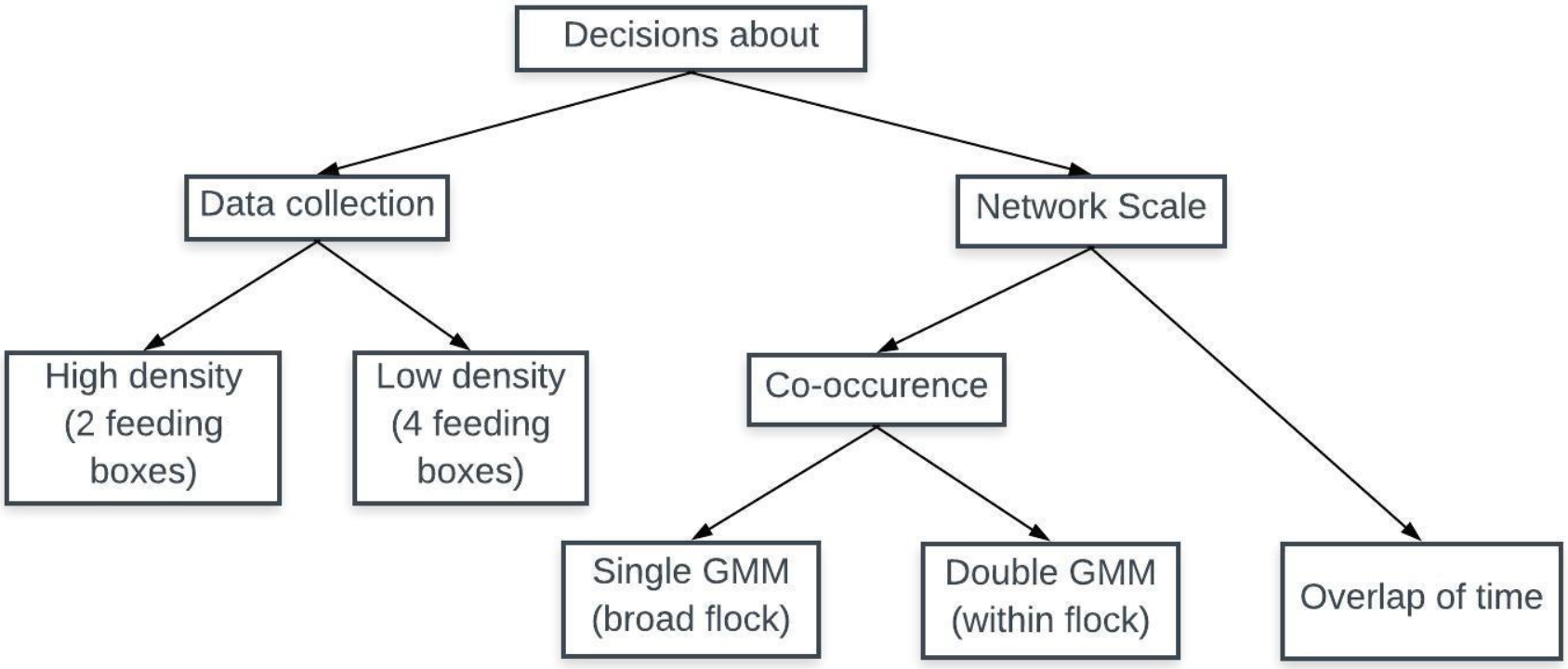
Flow diagram illustrating the steps for the two different aims of the study: comparing different methods for calculating edge weights and comparing different data collection setups.

#### Co-occurrence networks

##### Single GMM (broad flock)

We built networks using the rates of co-occurrence on the same so-called ‘gathering events’ as commonly done in other studies (e.g. Aplin et al. 2013). Gathering events were defined using a single run of the GMM (single GMM network) directly on the raw RFID feeders’ data which splits the temporal data in different foraging events based on peaks of activity on the feeding boxes (following Psorakis et al. 2015). We considered each feeding box as a different location to allow us to split the flock spatially in order to archive a greater resolution in detecting preferred association. We inferred the association strengths (edge weights) among colony members from their co-presence across all gathering events. We used the simple ratio index: the number of times that two individuals were in the same gathering event divided by the number of gathering events that contained at least one of the two individuals.

##### Double GMM (within flock)

Since our study species is colonial and highly gregarious, we believed that to differentiate the relationships among colony members we would need edges based on co-occurrences at a finer scale than what has traditionally been used for other species (the single GMM). Therefore, we used the Gaussian mixture model approach to define associations among individuals using a two-step procedure. Because the data from the feeders are quite discontinuous in this population (i.e. all individuals tend to visit foraging patches together and then all depart together in a very synchronised manner) we first detected the broader activity profile at the set of feeder boxes. We did this by grouping the individuals’ detections across all feeder boxes at a location into 1min blocks and used the GMM to extract the arrival and departure times of gathering events (see Fig. 2a). After this first step, we used the GMM again, but this time to detect waves of activity within each gathering event determined by the first GMM run. In the second run, we considered each feeder box (containing 4 RFID perches each) as a different location and used detections at a 1 second resolution. Considering each feeding box as a different location allowed us to split the data on the flock spatially, while running the GMMs within each gathering event allowed us to decrease the time scale and forced the GMM to split into shorter feeding bouts (Fig. 2b), thereby allowing the detection of within-flock spatial and social preferences. We inferred the association strengths among colony members from their co-presence across all feeding bouts generated from the second runs of the GMM (double GMM network). As with the single GMM approach we used the simple ratio index.

#### Time overlap networks

For the time overlap networks, we calculated the proportion of total feeding time during which two individuals were feeding simultaneously at the same feeding box (i.e. the time that birds spent feeding side-by-side). Here, edges were calculated by taking the sum of time that two individuals spent feeding at the same time at the feeding box divided by the sum of the total time that at least one of these two individuals were present at the feeder (which is also the simple ratio index, but more explicitly time-based rather than occurrence-based). This method aimed to define a stricter scale at which we consider that two individuals were associated, and represents the degree of tolerance to feed together. This method can be more meaningful for colonial and very gregarious species such as sociable weavers, since all members of the colony are often found foraging together and are already connected by colony membership, and since our interest is to find a sub-level of sociality within this colony structure.

### SNA analysis

We evaluated each network we produced by testing if they were significantly different from networks generated from randomizations of our data and if they generated patterns that reflect a biologically meaningful social aspect of this species: the assortment by members of the same breeding group. As we did not have a ground-truthed network against which to compare our networks, we instead used a meaningful social relationship—breeding group membership—to compare our different edge weight calculation methods, and ultimately evaluate which network performs best at detecting a sub-level of sociality within the colony that we expect to be present in the data.

Specifically, the utility of each network we generated (3 variants times 2 data collection methods) was evaluate according to two criteria:

1. The coefficient of variation (CV) in edge weights, to test which method would result in more differentiated networks. Low CV values represent a network in which individuals are equally connected, whereas a high CV value means that there are both strong and weak relationships detected. We used CV also to test if each network differed from networks generated from the randomization of the observation data.
2. Calculated weighted assortment coefficients (following Farine 2014) to determine which method better captured the expected preference to associate with members from the same breeding group. High values of assortment coefficients represent strong association between individuals of the same breeding group while low values represent no preference to associate with individuals of the same group. Not all individuals of the colony were attributed to a breeding group since not all breeding pairs managed to successfully reproduce during this breeding season, therefore for this analysis we used a subset of the network to include only individuals known to belong to a breeding group.

In order to test the statistical significance of the CV and the assortment coefficients, we compared the coefficients calculated from the observed networks with the same statistics calculated from 1000 random networks generated using permutations of the observed data (see Farine 2017). For the co-occurrence method, we generated random networks following the method first described by Bejder et al. (1998), using the R package asnipe (Farine 2013). Briefly, for the single GMM networks this method selects pairs of observations of individuals from different feeding bouts and then swaps these individuals. For the double GMM network the approach is similar, however pairs of observations of individuals are selected from the same broad flock visit (from the first run of the GMM) and at the same feeder, but from different gathering events (from the second run of the GMM). For the overlaps of time networks, we split the observed data by the gathering events defined by the first run of the GMM in the double GMM method and swapped the identity of the individuals within each gathering event. That is, we performed restricted node permutations (following Aplin et al. 2015b, but restricted by time rather than by space). By randomizing individuals’ detections events only within each gathering event, we aimed to keep constant, as much as possible, other factors besides social preferences that might contribute to the structure of the network (such as variation in individuals’ propensities to join flocks visiting feeders).

For all the 5 colonies we compared the CV and the assortment coefficients from the 3 different types of networks (singles GMM, double GMM and overlap of time). Additionally, for 2 of those 5 colonies we also compared the same 3 types of networks resulting from data collected using high and low density setups. This allowed us to test whether we could improve our networks not just in terms of edge definition but also regarding the design of data collection by changing the number of birds that can access food simultaneously. As illustrated in the diagram of Fig.3 the decisions about our method for constructing a suitable network for the sociable weavers were guided by both the setup design and the edge definition. Addressing these two questions might appear to be a sequential scheme, i.e. first looking at feeder saturation and after deciding if there was or not a significant improvement in using the 4 feeding boxes, addressing the scale problem (by comparing the different types of networks) or the other way (first the scale and then the feeder saturation). However, we did not address this as a sequential problem, since the two types of comparisons (comparisons of scale and comparisons of feeder saturation) are not easy to disentangle. In order to compare the high density with the low density setup we need a reliable network which can only be obtained by comparing the 3 types of networks. However, the 3 types of networks might be very different when using high feeding density setup but very similar when using a low density setup. For example, using the double GMM network with low density setup might be considered as good as a network from the high density setup using the overlap of time network. The results from both comparisons will guide the final decision on building the network and therefore must be presented and interpreted together and not in a sequential manner.

## RESULTS

All of the methods we used generated networks that were significantly different from random. From an edge definition perspective, the overlap of time method consistently generated networks with higher CV (Table 1) and higher values of assortment (Table 2). While the co-occurrence methods were able to detect the predicted positive assortment by breeding group in most colonies, the overlap of time method consistently produced considerably higher assortment coefficients. The single GMM co-occurrence method was able to generate well-differentiated networks, but performed worse with the assortment coefficients being close to zero (Table 2). These results suggest that the networks produced by the overlap of time method performed better at capturing a sub-level of sociality within the colony.

**Table 1.**
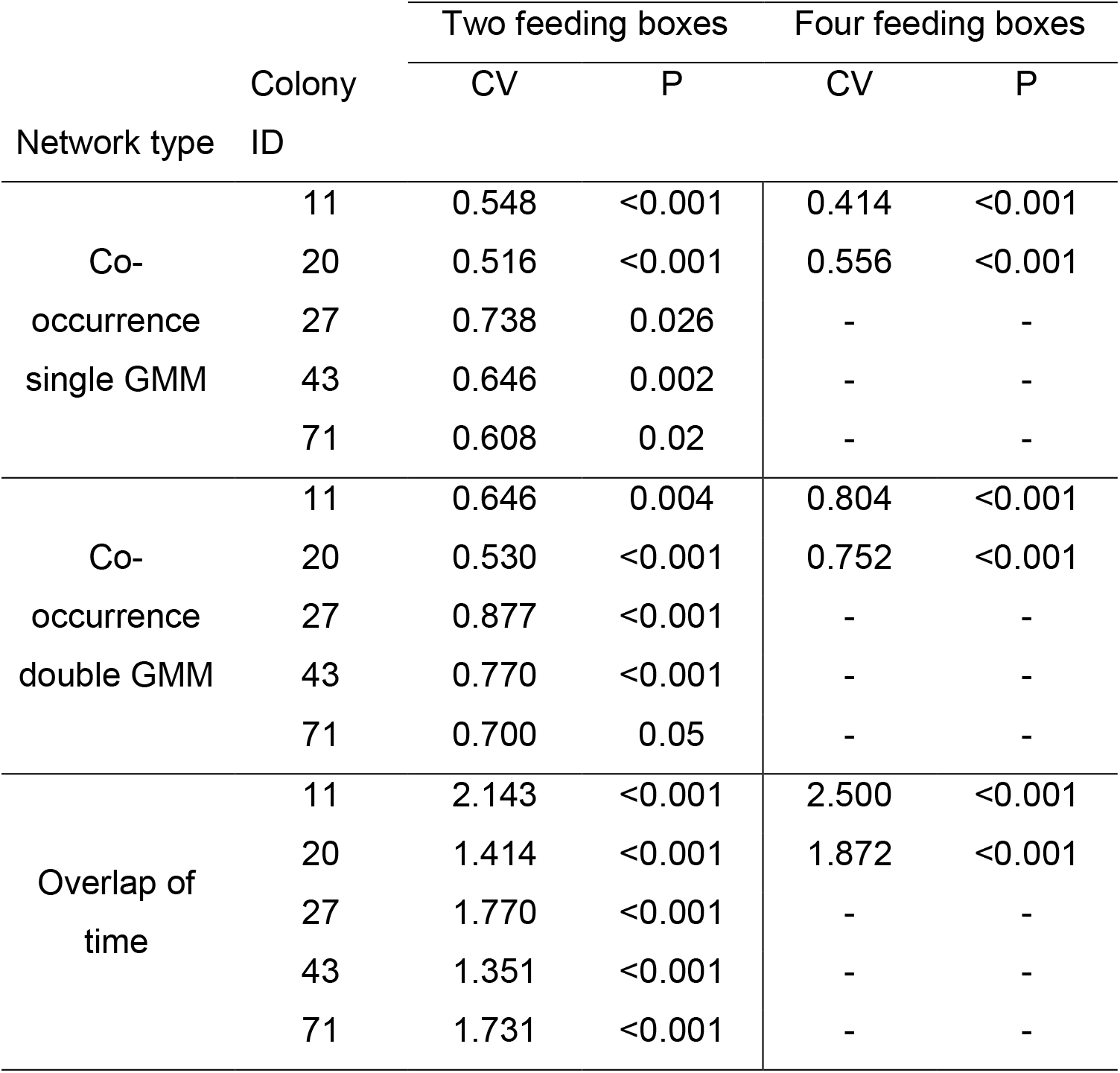
Comparison between the CVs of the three different types of networks obtained using a setup with two and four feeding boxes. Number of individuals per colony: colony 11: 34; colony 20: 27; colony 27: 38; colony 43: 27; colony 71: 59.

**Table 2.** Comparison between the assortment by breeding groups for the three different types of networks obtained using a setup with two and four feeding boxes. Number of individuals (number of groups) per colony: colony 11: 20 (8); colony 20: 10 (3); colony 27:20 (6); colony 43: 17 (5); colony 71: 19 (4).

| Network type | Colony ID | Two feeding boxes |  | Four feeding boxes |  |
| --- | --- | --- | --- | --- | --- |
|  |  | Assortment(SE) | P | Assortment(SE) | P |
| Co-occurrence single GMM | 11 | -0.005(0.026) | <0.001 | -0.020(0.027) | <0.001 |
|  | 20 | -0.063(0.086) | 0.14 | 0.053(0.094) | <0.001 |
|  | 27 | -0.017(0.028) | 0.002 | - | - |
|  | 43 | -0.013(0.013) | <0.001 | - | - |
|  | 71 | -0.005(0.039) | 0.018 | - | - |
| Co-occurrence double GMM | 11 | 0.018(0.029) | 0.004 | 0.092(0.045) | <0.001 |
|  | 20 | -0.022(0.088) | 0.12 | 0.232(0.105) | <0.001 |
|  | 27 | 0.009(0.032) | 0.052 | - | - |
|  | 43 | 0.049(0.041) | 0.012 | - | - |
|  | 71 | 0.012(0.042) | 0.002 | - | - |
| Overlap of time | 11 | 0.260(0.069) | <0.001 | 0.346(0.083) | <0.001 |
|  | 20 | 0.160(0.175) | <0.001 | 0.637(0.087) | <0.001 |
|  | 27 | 0.141(0.066) | <0.001 | - | - |
|  | 43 | 0.094(0.055) | <0.001 | - | - |
|  | 71 | 0.190(0.070) | <0.001 | - | - |

From a data collection methods perspective, using four boxes instead of two resulted in higher CVs and in higher assortment coefficients in both colonies (Table 1 and Table 2). In other words, using more feeding boxes at a given site resulted in greater power to discriminate between same breeding group associations for all the three types of networks. This effect was more pronounced in the co-occurrence method than in the overlap of time method.

## DISCUSSION

Using our empirical example, we have shown how different edge definition and experimental designs in the same context can result in different networks: some presenting a low coefficient of variation and thus a network in which individuals are more equally connected, and others with a higher CV, and thus a network containing a higher number of both stronger and weaker relationships. We also showed how knowledge about the study population can be used to help making decisions about the data collection design and determining how to calculate the strengths of social relationships. In the case of the sociable weaver, we showed that using the time that individuals spent together, rather than data on simpler co-occurrences, generates more meaningful networks. These networks had coefficients of variation in edge weights that were higher than the CVs of the networks generated from co-occurrences and clearly captured social structure. In fact, the assortment coefficients by breeding group were more than ten times higher in the time-based networks than in the networks generated from co-occurrences. Furthermore, we found that networks generated using more sampling opportunities (in this case a higher number of feeder boxes available simultaneously) were generally more meaningful, in line with the suggestions made in a recent methodological paper on sampling data for constructing networks (Davis, et al. 2018). More broadly, we were able to conduct meaningful statistical tests on the quality of the networks generated without influencing the evaluation of any future hypotheses we aim to test later, and therefore avoiding designs that resulted from multiple testing on a given hypotheses of interest.

We found that using more feeders simultaneously produced networks with higher assortment by breeding unit. Even though our analyses are based on only two colonies, the results suggest that using fewer feeders might reduce the ability for groups of preferred associates within a colony to forage together, or force them to forage with less preferred conspecifics. Alternatively, competition for access to the resource might imply that only more dominant individuals of each group can access the resource, while their subordinate preferred associates are prevented from associating freely. In either case, having fewer feeders means that birds could not clearly express their social preferences. In social network studies, the number of individuals that can be detected at the same time (or in a given time window) is rarely considered or reported (in this study 8 or 16 were used). Reporting the proportion of birds seen feeding could allow assessing if in some species, as it appears to be the case in the sociable weaver, using fewer detection points dilutes true social bonds, causing lower network resolution and potentially leading to less accurate associations. This issue becomes an important consideration for studies with limited budgets or researcher time. One will need to consider the trade-off between replication (sampling more individuals in total) and data quality (sampling more individuals simultaneously), as in our case, one setup requires twice as much equipment (or can sample at half the locations or revisit each location half as often). Simulation studies suggest that collecting more simultaneous data is generally preferable (Davis et al. 2018), because networks require many replicated observations of each possible pair of individuals in order to be robust (see Farine and Strandburg-Peshkin 2015).

We showed here that using the commonly used co-occurrence method was not the best option to reproduce the sociable weaver social structure. While using a more time resolved co-occurrence method (double GMM) resulted in a better network to capture assortment by breeding group, it still performed worse than a network based on the time that individuals spend in close proximity. This would be expected for a species such as the sociable weaver which forages in flocks always with the same individuals. While we found that a network definition based on the overlap in time provided the networks that best captured *a priori* knowledge of the study species’ social structure (i.e. the breeding group), it might not necessarily be the best method for all questions or study systems. It might be well suited to other colonial or highly gregarious animals living in very cohesive societies. We have shown the social behaviour of a given study species needs to be carefully considered when deciding how to collect data. Thus, new network studies should not simply implement the same methods from other systems, even if these were found to be most applicable there (such as using the single GMM method for fission-fusion flocks of great tits *Parus major*; Psorakis et al. 2015). Hence, for each study system, and potentially for each question, researchers should experiment with the best way to construct their networks, while avoiding trying the different methods on a given hypotheses of interest.

Our study also illustrates that different methods can generate different networks (see also Castles et al. 2014), and that *a priori* knowledge about the study species or population can be helpful in making decisions about which network to use. While different networks may be correlated (see Farine 2015), this does not mean they are all equally powerful at testing a hypothesis. However, when testing network quality, the choice of which *a priori* knowledge to use is also critical. For instance, failing to find a relationship between the overlap of time networks and a given individual attribute, such as personality, does not mean that individuals do not use information about these attributes when choosing their social partners. For example, in the European starlings (*Sturnus vulgaris*) the spread of a novel foraging task in a social group was predicted by a perching network but not by a foraging network, possibly as a result of a perching network better capturing social preferences than a foraging network in a captivity setup (Boogert et al. 2014b). Another example is the spread of foraging tasks from parents to offspring in most zebra finch families, where some juveniles seemed to explicitly learn from other adults instead of their parents (Farine et al. 2015). Collecting associations in different contexts or using different methods and testing these against different a priori knowledge should be an important part of most studies.

The crucial point to keep in mind, however, is that researchers should aim to make *a priori* decisions (even if some are inevitably arbitrary) about methods for collecting data and building networks that are independent of any later tests of hypotheses. Furthermore, reporting and discussing all the different pilot methods in published studies will be useful to others studies.

Here, we have tried to draw attention to the decisions that underlie social network analyses. Many recent papers provide guidance on how to construct networks (reviewed in Farine and Whitehead 2015). However, to our knowledge, little guidance is available about how to make system-specific decisions about data collection (e.g. number of individuals detected simultaneously) that can be critical to the results obtained. We show that integration of existing knowledge about the species’ social behaviour in making decisions can be a simple and very powerful way of informing which approach is the best one. The concepts we present using simple hypothesis testing to evaluate competing networks and help guide the process of building a network are easily generalised to other system. They go beyond breeding group membership (which is specific to cooperative breeders), foraging associations, RFID setups and co-occurrence *vs* time overlap used here, which we merely used as an empirical example. Any set of networks can be compared with a relevant biological metric regardless of the methods used.

In this paper, we provide a structured approach that can be used to make design decisions in network, or other, studies. In addition we also call for researchers to provide more information about the rationale leading to their decisions. In our case, we took advantage of the information obtained as a result of a long term project on cooperative breeding, which provided information on composition of breeding groups. In other projects this type of information might not be available or relevant, but others, such as the importance of mated pairs which are expected to share strong social bonds (see Boogert et al. 2014a; Firth et al. 2015) could be used. Finally, our study clearly highlights the need for data collection and analysis methods to be tailored to a given study system, as different approaches (all of which are valid and exist in the literature) produced quite different outcomes. We hope that once sufficient studies report their design process, as we have here, we will be able to identify some general guidelines for animal social network data collection and analysis.

## ACKNOWLEDGEMENTS

This study would have not been possible without the contribution of several people that helped with network data collection, to capture and monitor the reproduction of the birds at the study colonies: Rita Leal, Ryan Olinger, Sophie Lardy, Frédéric Simon, Annick Lucas, Nora Carlson, Antoine Grissot, Marion Devogel, Bárbara Reguera Alonso, Corie Jeal, Tendai Chino and Zingfa Wala. De Beers Consolidated Mines gave us permission to work at Benfontein Reserve. This study was supported by funding from the FitzPatrick Institute of African Ornithology (DST-NRF Centre of Excellence) at the University of Cape Town (South Africa), FCT (Fundação para a Ciência e a Tecnologia, Portugal) through grants IF/01411/2014/CP1256/CT0007 and PTDC/BIA-EVF/5249/2014 to R.C. and the French ANR (Project ANR-15-CE32-0012-02) to CD. ACF was funded by FCT SFRH/BD/122106/2016. DRF was funded by the Max Planck Society and the DFG Centre of Excellence 2117 “Centre for the Advanced Study of Collective Behaviour” (ID: 422037984) and CD by the CNRS. This research was conducted in the scope of the LIA ‘Biodiversity and Evolution’ (CNRS-CIBIO).

## Compliance with ethical standards

### Ethical approval

All experiments were conducted under permission of the Northern Cape Department of Tourism, Environment and Conservation (permit FAUNA1338/2017) and under the approval of the Ethics Committee of the University of Cape Town (2014/V2/RC). The procedures implemented in this work involved the capture, confinement, ringing, handling and blood sampling of adult birds and nestlings. All potential invasive procedures were conducted by experienced ringers. Adult birds were not kept for more than 3h and were released in small groups. Any bird showing signs of fatigue were kept in an aviary and released upon recovery.

